# Multiplexed Imaging Analysis of the Tumor-Immune Microenvironment Reveals Predictors of Outcome in Triple-Negative Breast Cancer

**DOI:** 10.1101/2021.01.06.425496

**Authors:** Aalok Patwa, Rikiya Yamashita, Jin Long, Michael Angelo, Leeat Keren, Daniel L. Rubin

## Abstract

Triple-negative breast cancer, the poorest-prognosis breast cancer subtype, lacks clinically approved biomarkers for patient risk stratification and treatment management. Prior literature has shown that interrogation of the tumor-immune microenvironment may be a promising approach for the discovery of methods to fill these gaps. Recently developed high-dimensional tissue imaging technology, such as multiplexed ion beam imaging, provide spatial context to protein expression in the microenvironment, allowing in-depth characterization of cellular processes. We demonstrate that profiling the functional proteins involved in cell-to-cell interactions in the microenvironment can predict recurrence and overall survival. We highlight the immunological relevance of the immunoregulatory proteins PD-1, PD-L1, IDO, and Lag3 by tying interactions involving them to recurrence and survival. Multivariate analysis reveals that our methods provide additional prognostic information compared to clinical variables. In this work, we present a computational pipeline for the examination of the tumor-immune microenvironment using multiplexed ion-beam imaging that produces interpretable results, and is generalizable to other cancer types.

## Introduction

Triple-negative breast cancer (TNBC) is a subtype of breast cancer that is negative for estrogen receptor, progesterone receptor, and human epidermal growth factor receptor 2. Representing an estimated 10-20% of breast cancers, it is characterized by aggressive behavior, including earlier onset, larger tumor size, and a more advanced grade^1,2^. TNBC is the subtype of breast cancer with the poorest prognosis^3^, having a lower chance of survival^4,5^ and higher risk of recurrence, especially within a short timeframe^6,7^. The absence of common breast cancer hormonal targets and high heterogeneity among TNBC tumors makes treatment management difficult, creating a need for more advanced interrogation of cellular processes within TNBC tumors^8^. Currently administered treatments, such as checkpoint inhibitors, only provide benefit to a small proportion of treated patients and are associated with high cost and toxicity^9^. Their effectiveness is limited, necessitating further interrogation of cancer-cell cues, factors in the tumor microenvironment, and host-related influences^10^. Currently, physicians are unable to separate patients with low risk of recurrence from patients with a high risk of recurrence, making it difficult to deescalate treatments for those who may not need it and pursue more aggressive treatments for those who do^11,12^. To risk-stratify for overall survival, the American Joint Committee on Cancer staging system is the most commonly used technique in clinical practice; it is based on variables such as tumor size, nodal status, and the presence of distant metastasis. However, its survival estimates vary considerably because other prognostically relevant factors are excluded^13^. There is a need to identify additional biomarkers of TNBC to aid prognosis ^14–16^. Identifying predictors of recurrence and survival in TNBC patients would allow improved patient stratification and targeted treatment plans, which would lead to better outcomes and spare patients from unnecessary aggressive therapies^17^.

The tumor-immune microenvironment (TIME) is a dynamic system comprising cancer cells, immune cells, and the surrounding extracellular matrix and vasculature^18^. The TIME is modulated by the expression and secretion of proteins that contribute to angiogenesis, immune suppression, and the coordination of the immune response^19^. Previous research has sought to discover the features of the TIME that are tumor-promoting or tumor-rejecting using transcriptomic and proteomic data^20–23^.

However, until recently, conventional histological techniques lacked the ability to measure the expression of a multitude of proteins at subcellular resolution while preserving spatial information^24,25^. Advancements in high-dimensional multiplexed imaging, such as multiplexed ion beam imaging (MIBI), have allowed for more direct interrogation of the TIME^26^ while boosting standardization and reproducibility of results^27^. MIBI uses secondary ion mass spectrometry to image antibodies tagged with isotopically pure elemental reporters^28^. It is compatible with formalin-fixed paraffin-embedded (FFPE) tissue samples, the foremost preservation method of solid tissue in routine clinical pathology. MIBI enables in-depth analysis of the TIME, measuring the expression of more than 40 proteins simultaneously while preserving spatial information^29^ and avoiding spectral overlap^30^ and autofluorescence^31^.

This study builds on the work of Keren et al.^25^, who found structure in the composition and spatial organization of the TIME. TIME architecture was broadly classified as immune cold, mixed, or compartmentalized, based on the amount of immune infiltration into the tumor. Immune architecture was associated with patient survival.

However, previous research did not test the association between single-cell features of the TIME and clinical outcomes such as recurrence/survival. Although previous work has identified macro-level features associated with survival, there is still a need to study more granular features of the TIME at subcellular resolution, such as the expression patterns of individual proteins, which can add prognostically relevant information^32^, and the characteristics of cell-to-cell interactions^33^.

In this work, we aim to uncover features of the TIME that are associated with recurrence and overall survival by analyzing MIBI scans of TNBC tissue^25,28^. The primary focus is to profile the proteins involved in cell-to-cell interactions and establish a link between the spatial organization of cells with varying expression patterns and clinical outcomes. We examine interactions involving functional proteins and immunoregulatory proteins in particular. As corollary aims, we demonstrate an association between protein co-expression patterns and recurrence/survival, examine proteins whose overall expression is associated with recurrence/survival, and test associations between immune composition and recurrence/survival.

## Results

### Patient Population

Our study examines 38 TNBC patients with no neoadjuvant treatment, a subset of the 41 patients examined by Keren et al.^25^ FFPE slides of breast tissue were taken from patients, scanned using MIBI, and subsequently segmented to demarcate cell boundaries.^25^ Patient data regarding age, tumor grade, stage, cancer site, and clinical outcome – recurrence and overall survival (OS) -- were also gathered (Table 1). We additionally gathered MIBI images of breast tissue of 8 healthy patients, a subset of the patients examined by Risom et al.^34^

**Table 1.** Patient cohort characteristics. SD refers to standard deviation. No. refers to the number, or count.

| Characteristic |  | All |
| --- | --- | --- |
| Patients, no. |  | 38 |
| Age, mean (SD) |  | 54 (15) |
| CS Tumor Size, mean (SD) |  | 38 (39) |
| Tumor Grade, no. (%) | 1 | 1 (3%) |
|  | 2 | 5 (13%) |
|  | 3 | 29 (76%) |
|  | 4 | 2 (5%) |
|  | Unknown | 1 (3%) |
| TNM Classification, no. (%) | 1 | 4 (11%) |
|  | 2 | 1 (3%) |
|  | 2A | 7 (19%) |
|  | 2B | 3 (8%) |
|  | 3 | 1 (3%) |
|  | 3A | 1 (3%) |
|  | 3B | 1 (3%) |
|  | 3C | 2 (5%) |
|  | Unknown | 17 (45%) |
| Cells per image, mean (SD) |  | 5006 (1527) |

### Dataset

MIBI scans produce images of protein expression from FFPE tissue, where each image has 44 channels; each channel conveys the expression of a certain marker on the tissue sample (Figure 1a). Cellular segmentations for both TNBC and healthy patients’ images were provided by Keren et al. and Risom et al., who utilized DeepCell, a deep learning technique for identifying individual cells from MIBI data^25,34,35^. Cell type assignment for TNBC patients’ images was also performed by Keren et al. through a hierarchical methodology (Figure 1b) (Methods).

**Figure 1.**
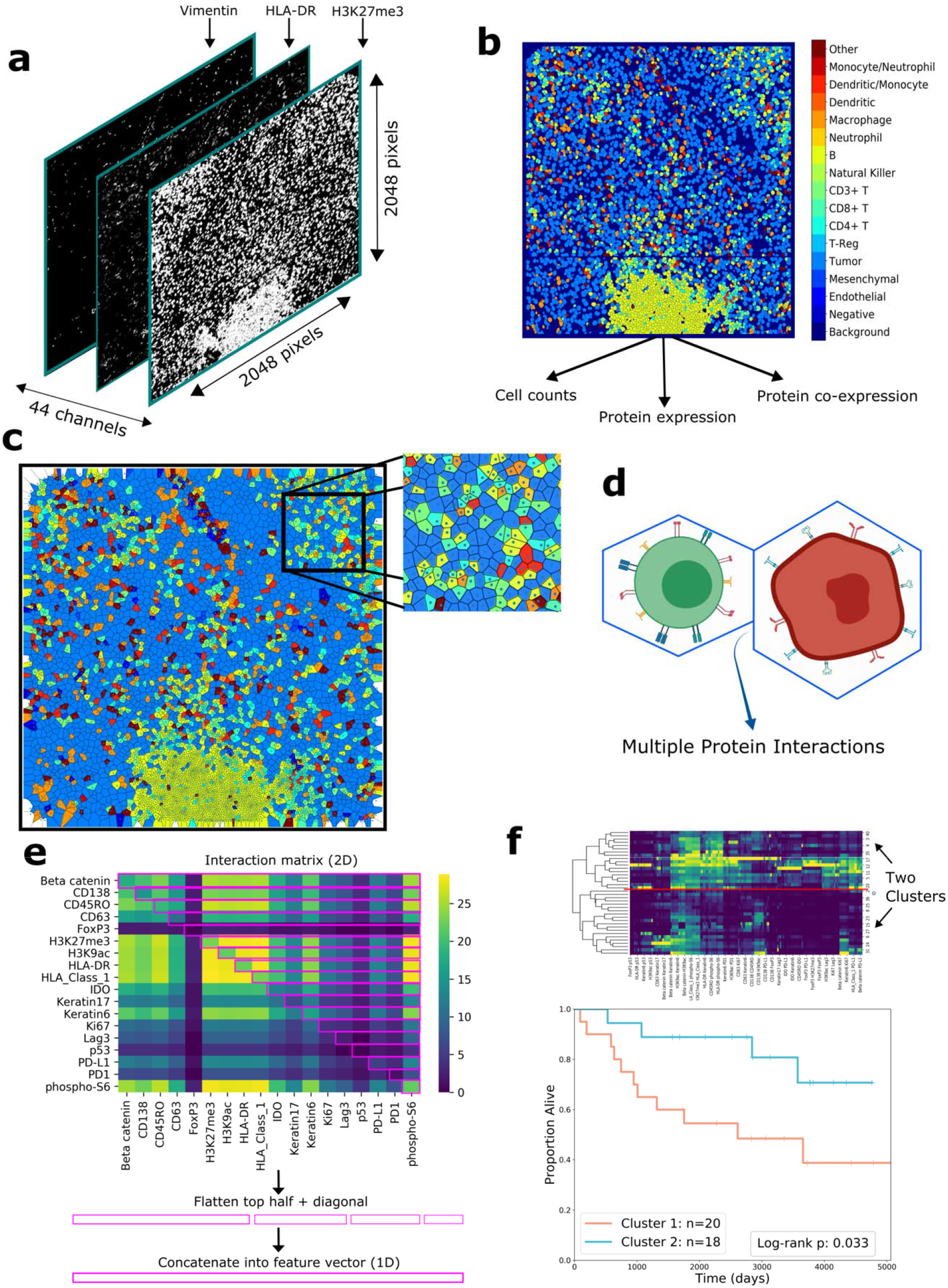
Overview of the computational pipeline. **a** Drawing of the layered structure of MIBI scans. Each MIBI image has dimensions of 2048×2048 pixels with 44 channels, where each channel represents expression for each protein; i.e., each pixel in the image at each channel conveys the concentration of that protein at that location. **b** Color-mapped image of cell segmentation performed on a MIBI image. The cell segmentation map has one channel with dimensions of 2048×2048. Each cell has its own cell type represented in colors referenced in the color bar on the right. From these cell segmentation maps and the original MIBI images, we extract cell counts, measure protein expression, and quantify co-expression. **c** Voronoi tessellation diagram of the cell segmentation map. Each polygon corresponds to a cell in the original segmentation, such that each point in the area of the polygon is closer to the centroid of the corresponding cell than any other cell. Each polygon borders a finite number of other polygons, simulating adjacencies between cells. **d** Using Voronoi diagrams, we analyze interactions between neighboring cells. **e** An interaction matrix is computed for each patient, with the entry at row A and column B representing the number of times a cell positive for protein A was adjacent to a cell positive for protein B (top). The top half triangle of the matrix, split across the diagonal, is selected, as shown with the purple rectangles. These rectangles are then flattened to form one feature vector, i.e., interaction features, for each patient. **f** Interaction features are used to cluster patients (top), and the two patient clusters are compared with regard to recurrence/survival using Kaplan-Meier plots and the log-rank test (bottom).

### Immune composition of cells is not associated with recurrence or survival

We examined whether the prevalence of certain cell populations in the TIME was associated with recurrence and survival. We measured the number of cells of each cell type in each patient and represented that number as a proportion of the total number of cells in that patient’s sample. We then performed univariate Cox regression and performed a two-sided t-test of the variable coefficient to determine whether each cell type’s prevalence was related to recurrence and overall survival.

After performing Benjamini-Hochberg correction to account for multiple comparisons^36^, there were no cell types whose coefficients were significant for either recurrence (Table 2a) or overall survival (Table 2b).

**Table 2a.** Immune composition Cox regression results for recurrence. There is no association between immune composition and recurrence in the cohort; no cell types had significant coefficients after correction.

| Cell Type | Coefficient | Hazard Ratio | Coefficient P | BH-Corrected FDR |
| --- | --- | --- | --- | --- |
| Monocyte/Neutrophil | -0.372 | 0.689 | 0.093 | 0.647 |
| CD8+ T | -0.072 | 0.93 | 0.154 | 0.647 |
| Macrophage | -0.053 | 0.949 | 0.182 | 0.647 |
| Mesenchyme | 0.037 | 1.038 | 0.183 | 0.647 |
| CD4+ T | -0.07 | 0.933 | 0.244 | 0.647 |
| Tumor | 0.011 | 1.011 | 0.286 | 0.647 |
| Natural Killer | -0.867 | 0.42 | 0.329 | 0.647 |
| Dendritic/Monocyte | -0.118 | 0.888 | 0.345 | 0.647 |
| Endothelial | 0.103 | 1.109 | 0.46 | 0.767 |
| B | -0.013 | 0.988 | 0.68 | 0.899 |
| Neutrophil | -0.046 | 0.955 | 0.682 | 0.899 |
| Dendritic | 0.022 | 1.022 | 0.739 | 0.899 |
| CD3+ T | 0.039 | 1.04 | 0.779 | 0.899 |
| Other | 0.004 | 1.004 | 0.942 | 0.971 |
| Regulatory T | -0.009 | 0.991 | 0.971 | 0.971 |

**Table 2b.** Immune composition Cox regression results for survival. There is no association between immune composition and survival in the cohort; no cell types had significant coefficients after correction.

| Cell Type | Coefficient | Hazard Ratio | Coefficient P | BH-Corrected FDR |
| --- | --- | --- | --- | --- |
| Dendritic/Monocyte | -0.292 | 0.747 | 0.076 | 0.435 |
| Tumor | 0.018 | 1.018 | 0.100 | 0.435 |
| Monocyte/Neutrophil | -0.331 | 0.718 | 0.121 | 0.435 |
| Other | -0.159 | 0.853 | 0.124 | 0.435 |
| Macrophage | -0.066 | 0.936 | 0.145 | 0.435 |
| Mesenchyme | -0.047 | 0.954 | 0.308 | 0.647 |
| CD3+ T | -0.169 | 0.845 | 0.315 | 0.647 |
| Dendritic | 0.048 | 1.05 | 0.41 | 0.647 |
| Neutrophil | -0.119 | 0.888 | 0.429 | 0.647 |
| Natural Killer | 0.357 | 1.43 | 0.431 | 0.647 |
| Endothelial | 0.102 | 1.107 | 0.505 | 0.689 |
| CD4+ T | -0.022 | 0.978 | 0.628 | 0.692 |
| B | -0.015 | 0.985 | 0.644 | 0.692 |
| Regulatory T | -0.129 | 0.879 | 0.646 | 0.692 |
| CD8+ T | 0.006 | 1.006 | 0.88 | 0.880 |

### Single-cell expression levels of functional proteins are not associated with recurrence or survival

We examined whether the expression of functional proteins in the cells of the tissue samples was associated with recurrence and survival (Figure 2a). We calculated the per-pixel expression levels of each protein in each patient. The histograms of expression for several proteins are shown in Figure 2b, and the histograms for all proteins are shown in Supplementary Figure 1. For this analysis, we included only functional proteins, which stand in contrast to proteins used solely for lineage assignment; their expression is modulated according to the functional state of the cell. Proteins whose expression had been implicated by previous literature as having an important role in tumor progression were designated “functional,” whereas those with less relevant roles were deemed “lineage.” We counted 18 markers in the “functional” category and 18 in the “lineage” category (Supplementary Table 1).

**Figure 2.**
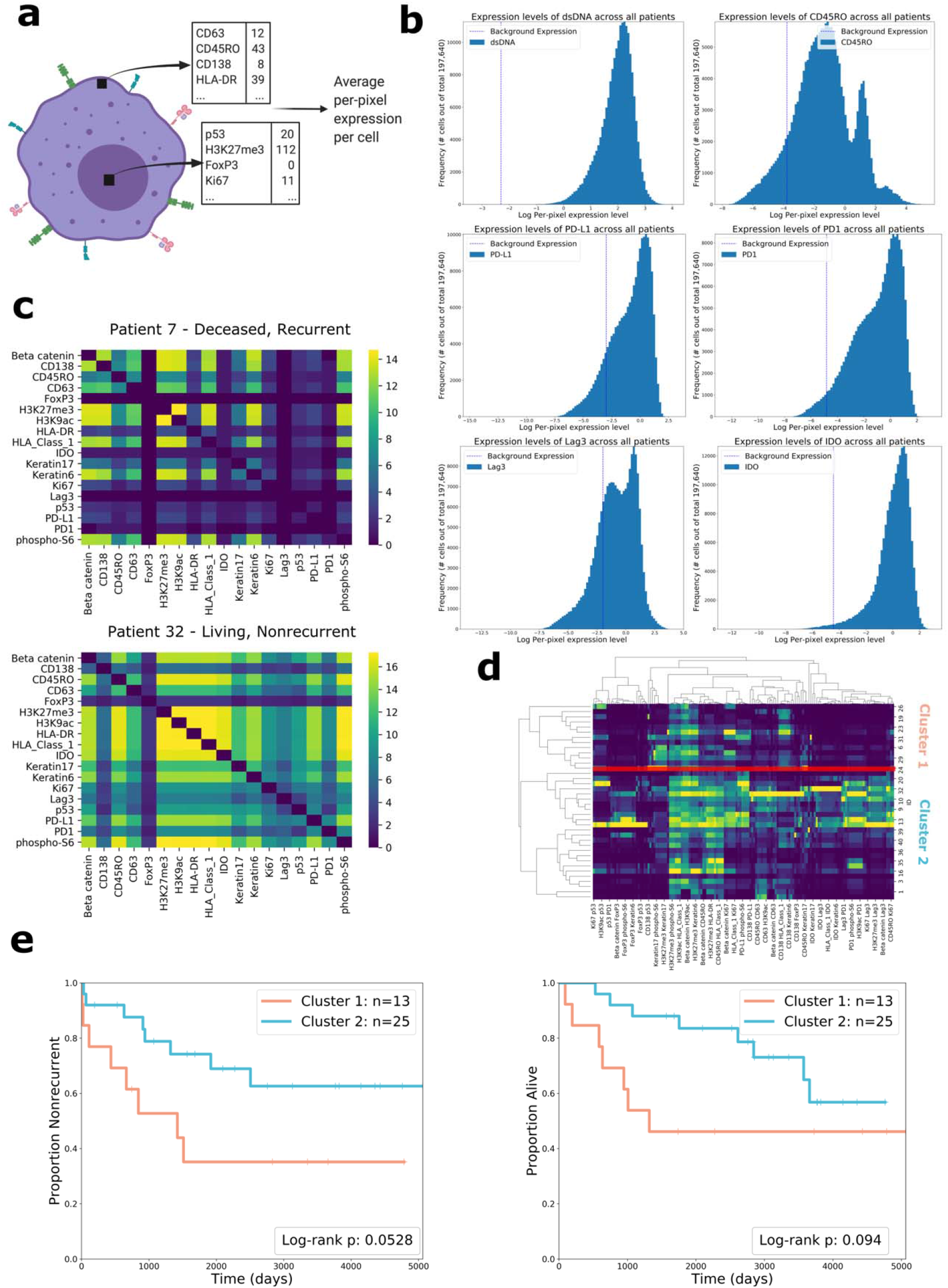
Quantification and analysis of protein expression. **a** Drawing showing how protein expression is calculated. The black squares each represent one pixel in the image. Expression levels are measured for each pixel in the cell and then summed across all pixels in the cell. The resulting number is divided by the size of the cell (in pixels), resulting in the average per-pixel expression level of the cell for each protein. **b** Histograms showing the distributions of log per-pixel expression levels for several relevant proteins. Per-pixel expression in the background channel (the positivity threshold) is shown with the vertical dotted line. **c** Heatmaps showing the cube root of coexpression of pairs of functional proteins in two different patients. The color bar also shows the cube root, so color value 16 indicates 16^3^ instances of co-expression. **d** Clustermap shows flattened features for all 38 patients. Two clusters were chosen from the dendrogram. The red line shows the way that the two clusters were separated. **e** Kaplan-Meier curves comparing clusters formed from co-expression features for recurrence (left) and overall survival (right). Two-sided log-rank test p-values are shown in the plot legend.

There were no functional proteins whose coefficients had significant p-values after Benjamini-Hochberg correction for either recurrence (Supplementary Table 2a) or overall survival (Supplementary Table 2b). Keratin6 (coefficient=0.025, HR=1.025, p=0.034) and HLA-DR (coefficient=-0.018, HR=0.982, p=0.045) were significantly associated with survival before correction. We placed Keratin6 and HLA-DR in a multivariate model to assess their relative prognostic relevance; Keratin6 remained significant (p=0.04), whereas HLA-DR did not (p=0.06). Expression of Keratin6 has been associated with poor survival outcome in previous work^37^, which our finding loosely corroborates. CD45RO (coefficient=-0.019, HR=0.981, p=0.051) was nearly significantly associated with recurrence before correction. CD45RO has previously been discussed in the literature for its role in anti-tumor immunity, especially with regards to its expression in memory T cells^38,39^. Our findings loosely corroborate this, as CD45RO expression was associated with favorable recurrence outcomes.

Within this cohort, the expression levels of functional proteins did not hold reliable prognostic relevance. As such, we decided to move away from macro-level interrogation of the TIME, opting to add spatial context to our analysis by quantifying protein co-expression and cell-to-cell interactions.

### Co-expression of functional proteins in patients’ cells is associated with recurrence and survival

We sought to develop a computational pipeline to test the association between localized coordination of immune activity and recurrence/survival. We calculated the number of times that pairs of functional proteins were co-expressed across all cells of a patient, summarizing this information in a “co-expression matrix.” (Figure 2c)

The co-expression matrices provide information regarding the phenotypes of the cells present in each patient, placing the expression of proteins in a single-cell context. We used the co-expression information as features to describe each patient. Patients were grouped by hierarchical clustering, and the tree was cut to form two patient clusters (Figure 2d). Our choice to select two clusters in this analysis, as well as all hierarchical clustering analyses, was motivated by silhouette score analysis^40^, which showed that division into two clusters would maximize inter-cluster dissimilarity (Supplementary Table 3). The recurrence/survival outcomes of the two patient clusters were compared using two-sided log-rank tests. They diverged according to recurrence (χ^2^(1, N=38) = 3.75, p=0.053), and survival (χ^2^(1, N=38) = 2.80, p=0.094) (Figure 2e). We also tested patient stratification when three clusters were chosen (Supplementary Figure 2). The log-rank test p-value for recurrence was 0.093 and 0.222 for survival.

We assessed the relative importance of individual co-expression features using random forest variable importance. The four most important co-expression features were CD45RO + H3K27me3 (score=0.822), CD45RO + H3K9ac (score=0.767), CD45RO + HLA Class 1 (score=0.646), and HLA-DR + IDO (score=0.604). These results show that calculating the coexpression of proteins, namely the combinations listed above, can aid patient stratification. CD45RO’s co-expression with HLA Class 1, an antigen used to promote cytotoxic T cell activation, is aligned with existing literature on melanoma^41^, and may evidence coordination between memory T cells and cytotoxic T cells in cancer.

### Cell-to-cell interactions contain prognostically relevant information

We examined cell-to-cell interactions by creating Voronoi tessellation diagrams out of the segmented MIBI images (Figure 1c). Voronoi diagrams have been used previously to define spatial organization and cellular morphology^31,42^. Each cell’s Voronoi polygon is created from the location of its centroid; its polygon will border some number of polygons from other cells^43^. These borders can be used to model cell-to-cell interactions (Figure 1d); cells whose polygons share a border can be considered adjacent (Figure 3a). Due to the geometry of the Voronoi tessellation algorithm, polygons will only border their immediate neighbors, which restricts the area of influence of a certain cell to the cells that are closest nearby.

**Figure 3.**
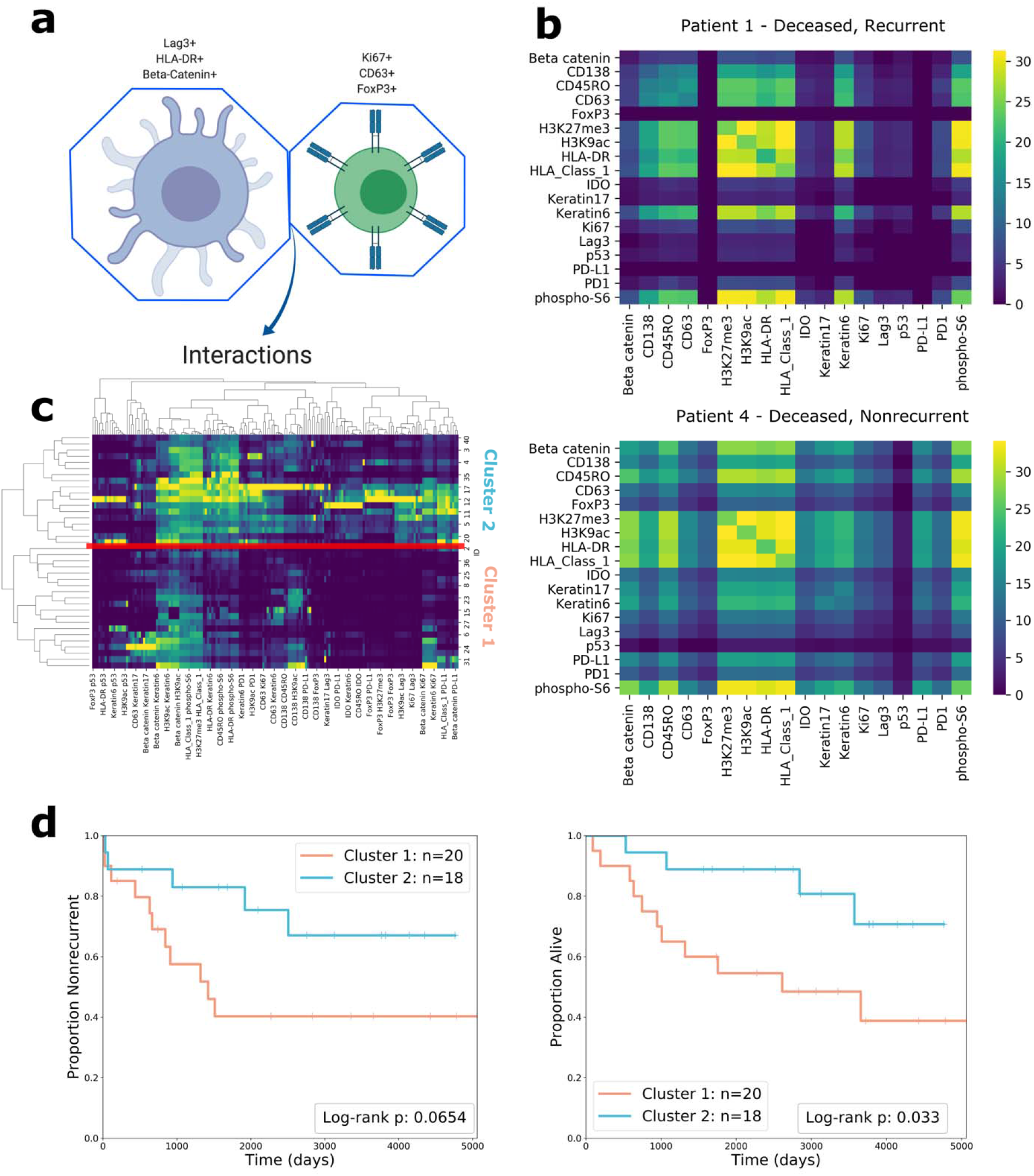
Analysis of cell-to-cell interactions. **a** Drawing shows how interactions are analyzed to find which combinations of proteins are involved in the interaction. The interaction is characterized by the adjacency of the tw**o** Voronoi polygons. Each cell involved in the interaction has a unique protein expression pattern, resulting in complex interactions. **b** Heatmaps showing the cube root of the number of interactions between pairs of functional proteins in two patients. The entry at row A and column B in the heatmap represents the cube root of the number of times that a cell positive for protein A was adjacent to a cell positive for protein B in that patient’s MIBI image. Pairs who had zero interactions are excluded from the plot. **c** Clustermap of patients’ functional protein interaction features. **d** Kaplan-Meier curves of recurrence (left) and overall survival (right) comparing clusters formed from interaction features. Two-sided log-rank test p-values are shown in the plot legend.

We created an interaction matrix for each patient to describe the characteristics of the patient’s cell-to-cell interactions by counting the number of times that specific pairs of proteins were involved in interactions. The entry in the matrix at row A and column B represents the number of times a cell positive for protein A was adjacent to a cell positive for protein B (Figure 3b).

Data for interactions involving functional proteins were used as features for hierarchical clustering, resulting in two clusters, with 17 patients in Cluster 1 and 21 patients in Cluster 2 (Figure 3c). The Kaplan-Meier curves comparing the clinical outcomes of the two patient clusters diverged according to recurrence (χ^2^(1, N=38) = 3.39, p=0.065) and diverged significantly according to survival (χ^2^(1, N=38) = 4.55, p=0.033) (Figure 3d).

Our method of quantifying cell-to-cell interactions reveals that the spatial proximity of functional proteins contains valuable prognostic information; the proteins involved in interactions can be used as features to cluster patients into groups with significantly different outcomes.

By contrast, quantifying interactions involving lineage proteins does not hold prognostic relevance. Hierarchical clustering on features of lineage protein interactions did not result in clusters that differed in recurrence and survival outcome significantly (Supplementary Figure 3).

A drawing comparing the clusters formed from clustering on functional protein interaction features to the morphology distinction performed by Keren et al. is shown in Supplementary Figure 4.

### Interactions involving immunoregulatory proteins predict recurrence and survival

We further examined a subset of functional proteins, the immunoregulatory proteins PD-1, PD-L1, IDO, and Lag3, which are in consideration as immunotherapy targets^9,44–48^. Prior research did not answer whether interactions involving these four proteins are associated with recurrence and survival, information that would be valuable in understanding their roles in TNBC progression.

To answer this question, we quantified spatial interactions between cells expressing immunoregulatory proteins, excluding all other proteins from the analysis (Figure 4a). We reasoned that interactions between cells positive for these proteins were associated with recurrence or survival, the result would point to the prognostic relevance of these proteins. Similar to previous analysis, the counts of interactions were used as features to cluster patients (Figure 4b). The Kaplan-Meier curves of the clusters formed from this analysis diverged significantly according to recurrence (χ^2^(1, N=38) = 7.60, p=0.0058) (Figure 4c). We also tested patient stratification when three clusters were chosen (Supplementary Figure 5). The three clusters diverged significantly with regards to recurrence (χ^2^(1, N=38) = 5.40, p=0.020), demonstrating that the efficacy of risk-stratification was robust to the number of clusters chosen.

**Figure 4.**
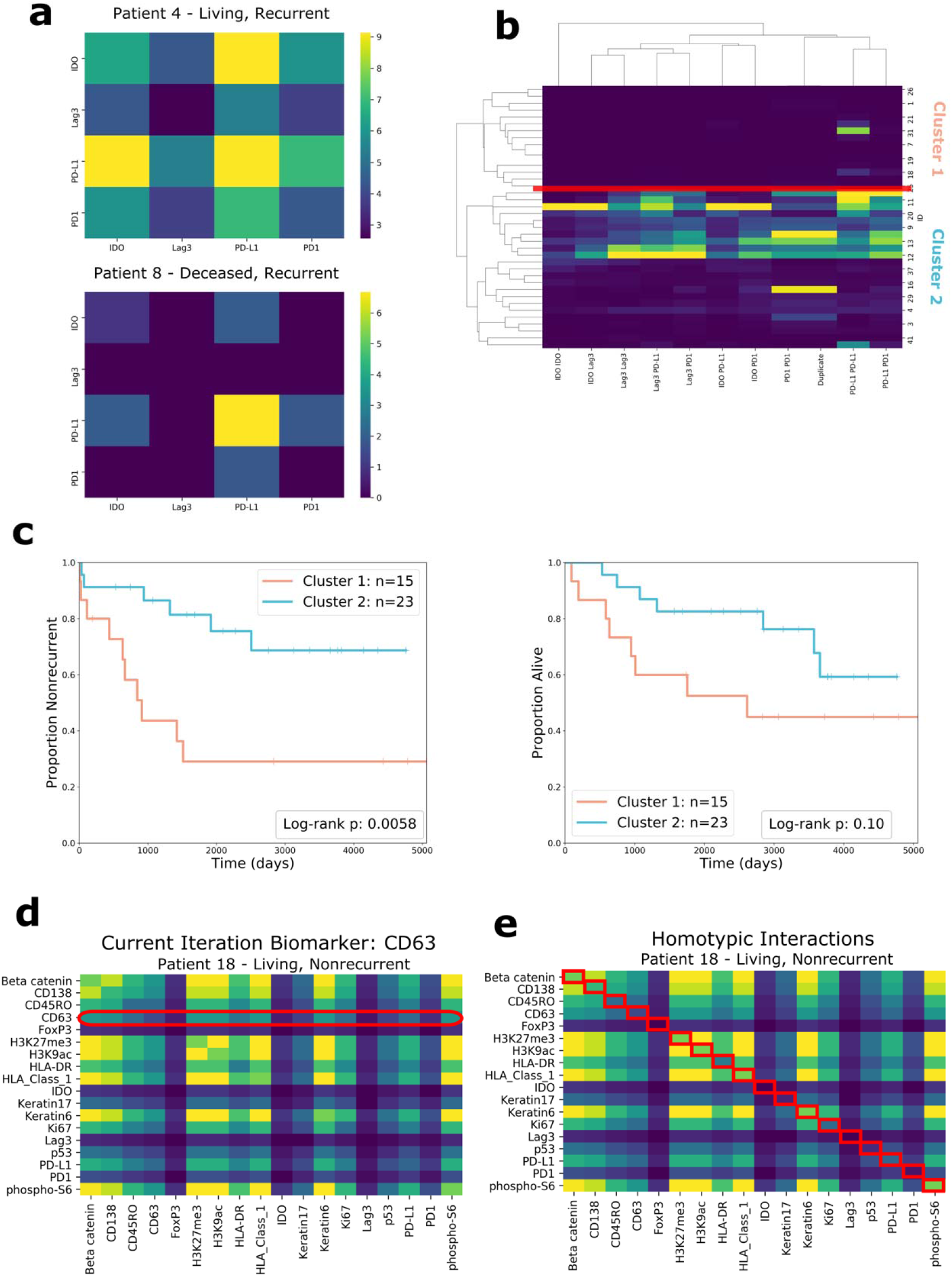
Analysis of subsets of interactions. **a** Heatmaps of the interaction matrices of immunoregulatory proteins (IDO, Lag3, PD-L1, PD-1) for two patients, whose outcomes are shown above the heatmaps. **b** Clustermap of patient’s immunoregulatory protein interaction features. The place at which the dendrogram was split is indicated with a red line. **c** Kaplan-Meier curves for recurrence (left) and survival (right) comparing clusters formed from immunoregulatory protein interactions. Two-sided log-rank test p-values are shown in the plot legends. **d** Diagram showing how the interactions of individual proteins are evaluated through ablation analysis one-at-a-time. The only interactions included as features are the ones that involve a specific protein. The diagram gives the example of CD63. **e** Diagram showing the set of homotypic interactions. As shown by the red boxes, only the entries in the diagonal are included as features.

### Ablation analyses reveal prognostically relevant groups of features

We further examined the cell-to-cell interaction data by performing multiple ablation analyses.

First, we examined individual functional proteins one-at-a-time, including only the interactions that involved this protein as features (Figure 4d). For example, when examining the interactions involving PD-1, we constructed feature vectors to include PD-1/Lag3 interactions, PD-1/Ki67 interactions, PD-1/PD-L1 interactions, and so on. There were several proteins whose interactions were significantly associated with recurrence: IDO (p=0.008), HLA Class 1 (p=0.011), H3K27me3 (p=0.011), and Beta Catenin (p=0.023). Phospho-S6’s interactions were significantly associated with survival (p=0.041).

We also examined “homotypic” interactions – interactions involving the same protein. Homotypic interactions are found in the diagonal of the interaction matrix – they represent the number of times in a patient that a cell positive for protein A was adjacent to a cell positive for protein A (Figure 4e). This information communicates the spatial proximity of cells with similar expression patterns. We used all of the homotypic interactions of functional proteins (the entire diagonal) as features for each patient and repeated the clustering analysis. The Kaplan-Meier curves diverged according to recurrence, χ^2^(1, N=38) = 3.43, p=0.064, and diverged significantly according to survival, χ^2^(1, N=38) = 4.90, p=0.027, indicating that the frequency of homotypic interactions is relevant information for survival prognosis.

We calculated the importance of interaction features by fitting a random forest model with interactions as predictors and cluster assignments as the response variable. Feature importance was scored using the mean decrease in Gini Index. The highest-importance feature was the Beta Catenin + CD45RO interaction feature (score=0.794), followed by CD45RO + HLA-DR (score=0.738), PD-1 + CD45RO (score=0.716), PD-1 + H3k27me3 (score=0.709), Lag3 + CD45RO (score=0.706), IDO + PD-1 (score=0.694), and Lag3 + PD-1 (score=0.647). CD45RO was present in 4 of the 7 most important interactions, PD-1 was present in 4, and Lag3 was present in 2. These results point to interactions involving these proteins as being particularly useful for patient stratification; they contributed the most to clustering, and the resulting clusters differed significantly in terms of recurrence and survival.

### Extracted features differ between healthy samples and TNBC samples

To confirm the validity of the features we extracted, we tested whether they differed between healthy tissue samples and TNBC tissue samples. The healthy tissue used in our analysis came from a different study, which profiled a different set of markers. There were 6 proteins common between the healthy images and TNBC images: FoxP3, IDO, Ki67, PD-1, PD-L1, and phospho-S6.

We calculated expression levels for the healthy tissue and compared them against the TNBC tissue using a two-sided Wilcoxon rank-sum test. Five out of the six proteins were significantly different across tissue: FoxP3 (W=2.06, p=0.040), Ki67 (W=3.06, p=0.022), PD-1 (W=3.622, p=0.0003), PD-L1 (W=2.42, p=0.020), and phospho-S6 (W=4.00, p=6.35e-05). Bar plots comparing healthy and TNBC tissue for each protein are shown in Supplementary Figure 6a.

We also validated our method of profiling cell-to-cell interactions on the healthy tissue by subjecting it to our computational pipeline and testing whether the cell-to-cell interaction features of healthy tissue would be different from TNBC tissue. We reduced the interaction features for healthy and TNBC patients to two dimensions using UMAP (Uniform Manifold Approximation and Projection)^49^ and plotted the reduced features for visualization. The resulting scatterplot showed separation between healthy and TNBC tissue (Supplementary Figure 6b).

These results suggest that our computational pipeline succeeded in extracting tumor-specific single-cell spatial features that are prognostic for recurrence and overall survival. Additionally, they demonstrate that our computational pipeline is applicable to a variety of MIBI datasets, as we applied the same methods to the two distinct datasets.

### Multivariate analysis reveals features with independent prognostic relevance for recurrence and survival

To assess the prognostic importance of the features we identified, we fitted three multivariate Cox regression models, each of which included one of the cluster variables, two clinical variables (grade and age), and the immune architecture distinction described by Keren et al. We obtained coefficients and hazard ratios to determine whether the cluster variables added prognostic information.

Both of the clusters formed from cell-to-cell interaction features contained additional prognostic information for at least one clinical outcome. The immunoregulatory proteins interaction cluster contained independent prognostic information for recurrence (coefficient=-1.32, HR=0.27, p=0.02). The functional proteins interaction cluster contained independent prognostic information for survival (coefficient=-1.24, HR=0.29, p=0.04). These results suggest that our computational pipeline was able to extract additional prognostically relevant features and use them to risk-stratify patients.

Next, we assessed the relative prognostic relevance between each of the cluster variables. To do this, we fit random forests with six predictors: the three cluster variables, two clinical variables (tumor grade and age), and the immune architecture distinction defined by Keren et al. We then measured variable importance by calculating SHAP (Shapley Additive Explanations) values^50^ and overall goodness-of-fit using Harrell’s c-index^51^.

The random forest analysis corroborated our results from the multivariate Cox regression analysis. The immunoregulatory protein interactions cluster was the most relevant feature for recurrence (Figure 5a), and the functional protein interactions cluster was the most relevant feature for survival (Figure 5b). These features were more important than tumor grade, age, and tumor architecture. The c-index for the recurrence model was 0.718, and the c-index for the survival model was 0.731, indicating a good fit.

**Figure 5.**
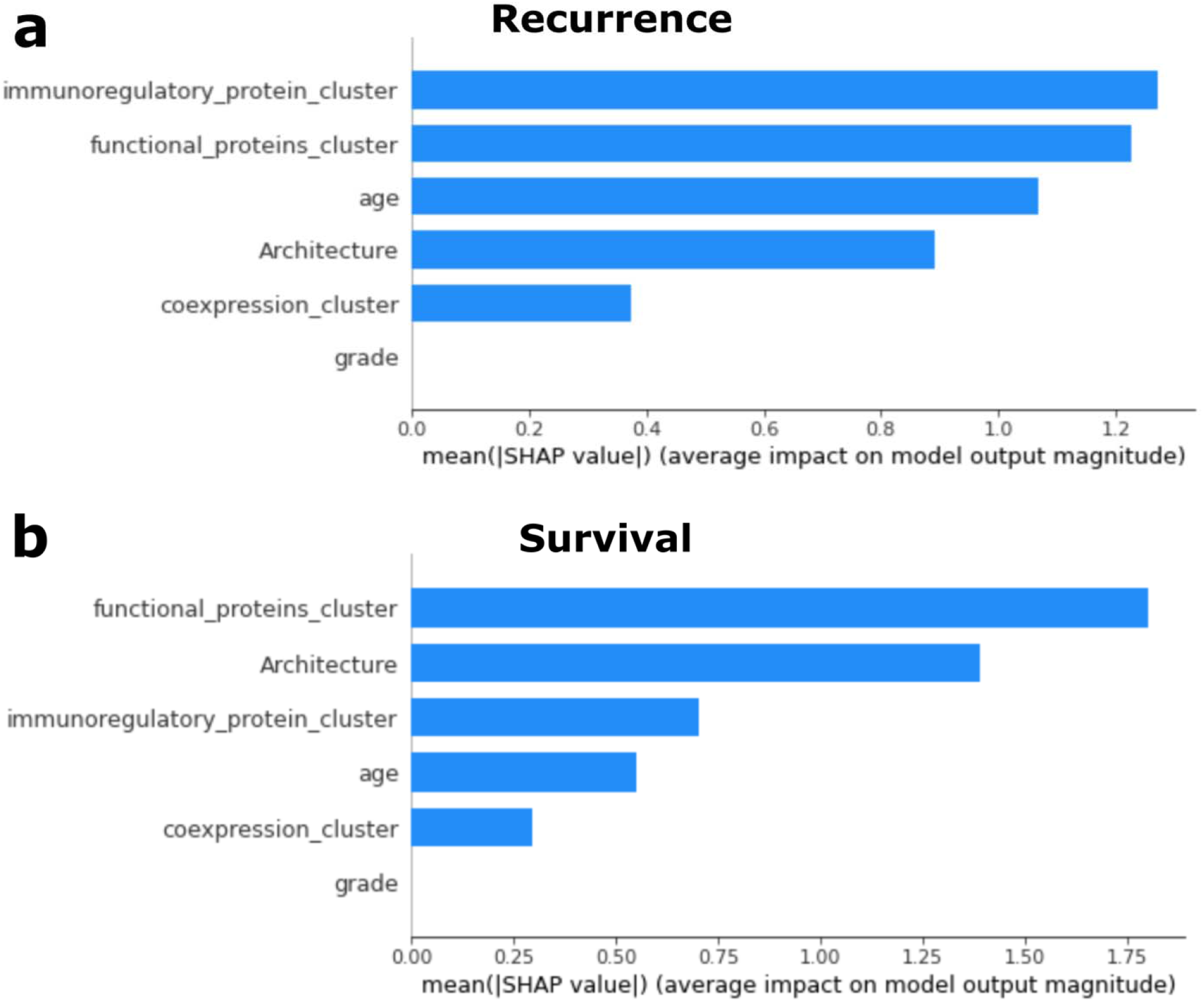
Random forest variable importance. **a** Bar plot showing the mean SHAP value for each variable in a random forest predicting recurrence. SHAP (Shapley Additive Explanations) values are a measure of variable importance that quantify how the expected model prediction would change when conditioning on a certain variable. They are more aligned with human intuition than other feature attribution methods. **b** Bar plot showing the mean SHAP value for each variable in a random forest predicting survival.

## Discussion

TNBC is the most aggressive breast cancer subtype, with a higher risk of recurrence and lower probability of survival. It lacks clinically approved biomarkers for patient risk stratification, making treatment planning and management difficult. Previous research involving TNBC and multiplexed imaging did not analyze the prognostic relevance of protein co-expression patterns and cell-to-cell interactions. In this study, we aimed to examine the association between these features and recurrence/survival in TNBC patients by constructing a computational pipeline for the analysis of MIBI.

Our contributions are threefold. First, we identify possible predictors of recurrence and overall survival in TNBC, demonstrating that the information contained within cell-to-cell interactions and protein co-expression patterns can aid patient stratification and therapeutic design, as proven through evaluation of patient groups, statistical tests, and predictive modeling. Second, we demonstrate that the immune composition of the TIME does not always hold prognostic relevance, and should therefore be examined with caution. Third, we present a computational pipeline for the interrogation of the TIME that produces interpretable and conclusive results, making it potentially viable in a clinical setting.

Our primary focus was to examine cell-to-cell interactions in the TIME for the purpose of patient risk stratification and treatment management. Our findings show that the type and number of cell-to-cell interactions involving functional proteins quantified by our pipeline were associated with both recurrence and survival in our cohort, and could possibly serve as a tool for prognosis. The two most important interaction pairs were CD45RO + Beta Catenin and CD45RO + HLA-DR, a finding that corroborates underlying biology. CD45RO marks memory T cells, which have been shown to mediate anti-tumor immunity^38,39^. Beta Catenin is expressed on tumor cells primarily^52,53^, so its interaction with CD45RO evidences the anti-tumor actions of memory T cells. HLA-DR is expressed on antigen-presenting cells^54^, so its interaction with CD45RO+ cells evidences coordination between different immune cells to suppress tumor growth. The biology behind these interactions demonstrates that computational analysis of cell-to-cell interactions can elucidate immunological mechanisms playing a role in patients’ tumors. The biological significance of the proteins involved in these interactions could be further investigated through biological analysis of animals or clinical trials. Further, we found that “homotypic” interactions – interactions involving the same protein – hold predictive power. This finding indicates a coordinated immune response characterized by the localized enrichment of functional proteins.

Multivariate analysis revealed that the interactions of functional proteins contained independent prognostic information for survival, even when compared to clinical variables like tumor grade, age, and the architecture distinction determined by Keren et al., pointing to the potential efficacy of this technique for patient stratification and treatment management of TNBC. We also profiled the cell-to-cell interactions of healthy tissue. The interaction features of the healthy tissue were distinct from the interaction features of TNBC tissue in our cohort, indicating that profiling cell-to-cell interactions may also be able to convey information regarding the pathological state of tissue. However, further research with a larger sample size is necessary to confirm this.

We further analyzed four immunoregulatory proteins – IDO, Lag3, PD-1, and PD-L1 – which are in consideration as immunotherapy targets^9,44–48^. We found that the expression profiles of these proteins were prognostically relevant, suggesting that these proteins can play a role in modulating tumor progression. A host of literature has described the importance of these proteins in TIME processes^9,32,44,55^, but only a small subset of such literature examines them in the context of paired cellular interactions. Interestingly, the individual expression levels of these proteins were not prognostically relevant; after Benjamini-Hochberg correction, none of the proteins had expression levels significantly associated with recurrence or survival. However, the cluster variable formed from their interactions was highly predictive of recurrence, as shown by multivariate analysis. Profiling the cell-to-cell interactions involving immunoregulatory proteins revealed independent prognostic information when compared to tumor grade, age, and the architecture distinction determined by Keren et al.

Our methods differ from the previous analysis of these data in several ways. Keren et al. calculated interaction matrices by defining a distance of 39 micrometers to establish adjacent cells^25^; however, the features within these interaction matrices did not result in patient clusters that differed significantly with respect to clinical outcome. This may suggest that using a set distance for adjacency is of insufficient spatial resolution to differentiate microenvironments. Our analysis also used a much lower threshold for cell protein positivity. This lower threshold may have improved the detection of important interactions. Voronoi diagrams and Delaunay triangulation have been used previously to define and examine cellular neighborhoods in colorectal cancer^31,56^. In contrast, we use Voronoi diagrams to examine protein expression in pairwise cellular interactions, rather than larger neighborhoods. We then use these pairwise cellular interactions to explain higher levels of abstraction, building interaction matrices to summarize patients’ TIME overall.

We found that the co-expression profile of functional proteins in patients’ cells is associated with recurrence and survival. The four most important co-expression pairs were CD45RO + H3K27me3, CD45RO + H3K9ac, CD45RO + HLA Class 1, and HLA-DR + IDO. These results point to highly specific cellular phenotypes, a trademark of a complex TIME^25,57,58^. Our computational pipeline presents an efficient, interpretable way to identify co-expression patterns and use them to risk-stratify patients.

Our methodology allowed for analysis of the cell types present in the TIME as a whole, providing a macro-level view of immune coordination. Our findings indicate that caution should be exercised when using the immune composition as a biomarker in clinical settings. After Benjamini-Hochberg adjustment, there were no cell types with significant prognostic value. This does not corroborate existing literature regarding the prognostic relevance of certain cell types, including the monocyte/neutrophil cell type^59^, the dendritic cell/monocyte cell type^60^, natural killer cells^61,62^, CD8+ T cells^27,63,64^, macrophages^65^, B cells^66^, CD4+ T cells^67,68^, and CD3+ T cells^69^.

The subcellular resolution achieved by MIBI allowed us to quantify the expression of individual molecules on a single-cell basis. After Benjamini-Hochberg correction, there were no proteins significantly associated with either recurrence or survival. Keratin6 and HLA-DR were associated with survival before adjustment, and Keratin6 remained significantly associated with survival when placed in a multivariate model with HLA-DR. This aligns with some existing literature. CD45RO was almost significantly associated with recurrence before adjustment, but its prognostic relevance was more clearly highlighted through its cell-to-cell interactions and co-expression patterns. This suggests that adding spatial context to the TIME can reveal otherwise hidden prognostically relevant information, a potential benefit of our developed computational pipeline for MIBI analysis.

A limitation of our work is that our results are derived from a sample of 38 TNBC patients that were treated at Stanford hospital from 2002 to 2015 – further work is needed to validate these results on a larger cohort of patients. Although it was known that the patients had not undergone neoadjuvant treatment, further data regarding treatments pursued was not available; future research is necessary to examine associations across treatment types. In addition, this study was retrospective and performed with patients at a single institution. Our cell type classifications were found computationally, derived only from the expression of molecules that were a part of our chosen assay – future work should repeat this analysis using other biologically relevant molecules.

Nonetheless, this study presents a computational pipeline for the robust interrogation of multiple features of the TIME. We demonstrate the potential for cell-to-cell interactions and protein co-expression to improve prognosis and patient stratification. We found several statistically significant results within a limited cohort, suggesting that they may have large effect sizes and merit further exploration. Our methods produce interpretable results, which may make them beneficial in therapeutic design^70^. Furthermore, they can be applied to other cancer types, as they are generalizable to any MIBI scan.

## Methods

### Patient Population and Dataset

Our study examined 38 TNBC patients who were treated at Stanford Hospital from 2002-2015, a subset of the cohort examined by Keren et al.^25^ None of the 38 patients had undergone neoadjuvant treatment. Although the original cohort contained 41 TNBC patients, 3 of the patients were unusable for our analysis. Patients 22 and 38 lacked recurrence outcomes, and Patient 30’s images were corrupted. These patients had no special type, with estrogen receptor and progesterone receptor positivity less than 1% and HER2 unamplified. 1mm cores were taken from each patient and H&E stained. All samples were then stained with an antibody mix and scanned using MIBI-TOF. A computational pipeline converted the output of MIBI-TOF into images.

The dataset included two separate sets of 2048 x 2048 pixel images, representing a region of 800^2^ square micrometers. The first set of images are 44-channel TIFFs that represent protein expression levels, where each patient has one TIFF. Each channel in the TIFF corresponds to one of the 44 molecules profiled in the study. Of the 44 molecules, 36 were biological macromolecules, such as double-stranded DNA or IDO, and 8 were elemental reporters. Each pixel in the image has a value representing the expression of the protein in that location. The second image set contained 38 grayscale segmentations of cells in the patient’s sample. Patient data regarding age, tumor grade, stage, and recurrence and survival outcomes were also gathered.

The cellular segmentation and cell lineage clustering were performed by Keren et al.^25^ To perform nuclear segmentation, the authors utilized DeepCell, a deep learning-based method for segmentation of MIBI data^35^. The model was trained using manual segmentations of patients 1 and 2 and run on the images of all patients. Cell boundaries were defined as a 3-pixel radial expansion around the nuclei. Cells were clustered hierarchically. First, they were clustered into “Immune” and “non-immune” using the expression levels of CD45, FoxP3, CD4, CD8, CD3, CD20, CD16, CD68, MPO, HLA-DR, Pan-Keratin, Keratin17, Keratin6, p53, Beta catenin, and EGFR. Non-immune cells were clustered into Epithelial, Mesenchyme, Endothelial, and Unidentified using Vimentin, SMA, CD31, Beta-catenin, EGFR, Keratin16, Keratin6, and Pan-Keratin. Immune cells were further clustered into 12 groups (Figure 1b) using CD4, CD16, CD56, CD209, CD11c, CD68, CD8, CD3, CD20, HLA-DR, CD11b, MPO, and FoxP3.

We additionally gathered 8 MIBI images of breast tissue of healthy patients. These MIBI images were originally collected for a different study, and they profiled a different set of markers. Six proteins overlapped between the TNBC samples and healthy samples: FoxP3, IDO, Ki67, PD-1, PD-L1, and phospho-S6.

The cellular segmentation of MIBI images of healthy patients was performed by Risom et al.^34^ using DeepCell. Two distinct segmentations were performed. The first applied a three-pixel radial expansion and a stringent threshold for splitting cells. The second applied a one-pixel radial expansion and a lenient threshold. A post-processing step gave preference to the lenient threshold when the two segmentations were combined.

The authors complied with all ethical regulations involving human clinical data. Informed consent was obtained for all participants by previous studies. The study protocol was approved by the Stanford University Institutional Review Board.

### Analysis of Cell Prevalence

We examined whether the cellular composition of the TIME was associated with recurrence and survival. We quantified the number of cells of each cell type in each patient. To isolate specific cell types at a time, we created binary masks of each grayscale value to isolate each cell type. Then, we found the number of connected components in each mask, which provided the number of cells of each cell type. After noticing a large variation in the total number of cells per patient, we divided each patient’s cell type count by the total number of cells in their TIME to control for this lurking variable. Univariate Cox regression was then performed for each cell type to assess its association with recurrence and survival. Regression coefficients were examined using twosided t-tests. P-values were adjusted using the Benjamini-Hochberg method^36^.

### Single-Cell Protein Expression

We examined whether the expression levels of functional proteins within patients’ TIME were associated with recurrence and survival. For this analysis, we analyzed functional proteins, which modulate the activity of the cells in the TIME. These proteins stand in contrast to proteins used solely for lineage assignment; their expression is related to the functional state of the cell.

We labeled connected components and created a binary mask of each component to isolate the space taken up by each cell. We then applied this mask to the MIBI protein expression images, summing the value in each channel of the TIFF for each pixel in the mask. This created a 44-length vector of protein expression per cell. We realized that per-cell expression levels are dependent on cell size, so we divided the 44-length expression vector by the size (in pixels) of the cell. This left a 44-length vector representing average per-pixel protein expression for a certain cell.

We calculated protein positivity thresholds from the expression levels of the image background, which lacks cells and therefore can act as a negative control. We calculated total protein expression in all background pixels in all patients and divided these values by the total number of background pixels across all patients (~67,000,000 pixels). We used each protein’s threshold value to determine whether a cell was positive for a certain protein. Then, we counted the number of cells in each patient that were positive for each protein and divided the number by the total number of cells in the patient. The result was the proportion of cells in the patient that were positive for this protein. For example, 100% of a patient’s cells would be positive for DNA, but only 30% might be positive for PD-1. Univariate Cox regression was used to determine the association between protein expression proportions and clinical outcomes. Regression coefficients were examined using two-sided t-tests. P-values were adjusted using the Benjamini-Hochberg method.

### Functional Protein Co-expression

The co-expression of functional proteins in a single cell reveals functional status and immune coordination. We assessed the association between co-expression of functional proteins and recurrence and survival. We had previously measured single-cell protein expression and determined a threshold to designate cells as “positive” for each protein. We defined coexpression as an instance in which an individual cell is positive for a pair of proteins. For example, if a particular cell is positive for IDO, Lag3, and PD-1, it would have 3 instances of coexpression: IDO/Lag3, IDO/PD-1, and Lag3/PD-1. We constructed an 18×18 co-expression matrix for each patient to summarize the number of cells in the patient that co-expressed each pair of proteins. Because these matrices were symmetrical (a co-expression of IDO/Lag3 is the same as a co-expression of Lag3/IDO), we divided the matrix in two across the diagonal and flattened the top half to create a feature vector for each patient. To control for lurking variables, we standard scaled features across patients. We then performed hierarchical clustering to segment patients according to these features.

Silhouette analysis^40^ revealed that choosing two clusters would lead to the optimal segmentation, so we cut the dendrogram into two distinct clusters and compared the two groups using a twosided log-rank test. Then, to assess the importance of individual co-expression features, we fit a random forest with all of the co-expression pairs as predictors and the cluster assignment as the response. We assessed variable importance using a mean decrease in the Gini index.

### Voronoi Tessellation

Analyzing cell-to-cell interactions requires a method of defining cell adjacencies. We used Voronoi tessellation diagrams to model cellular adjacencies within the TIME. Voronoi tessellation divides a planar space into a number of regions such that each point in the plane has its own region in the tessellation^43^. The sides of each Voronoi polygon are constructed to bisect two input points. Therefore, each line segment in the Voronoi tessellation represents the borders between two input points. Voronoi diagrams have been applied to single-cell imaging technology in the past, specifically for visualizing the spatial organization of colorectal cancer cells^31,42^. Due to the geometry of the Voronoi tessellation algorithm, polygons will only border their immediate neighbors.

We labeled connected components from the cell segmentation images to find each cell’s centroid, which was then used to create Voronoi diagrams for each centroid. Therefore, every cell in the original cell segmentation images has a corresponding Voronoi diagram. We considered cells with bordering Voronoi regions to be adjacent, and therefore interacting. This created a reliable foundation for upstream analysis.

### Cell-to-cell Interaction Analysis

We used the borders created by Voronoi diagrams to iterate over all cell adjacencies in each MIBI image. Each adjacency represented an individual interaction between two cells. We constructed two lists: List 1 contained the names of the proteins that Cell 1 was positive for, and List 2 contained the names of the proteins that Cell 2 was positive for. We took the Cartesian product of the two lists to find all of the combinations of proteins present in this interaction. For example, if Cell 1 was positive for PD-L1 and Lag3, and Cell 2 was positive for PD-1 and IDO, then we would count the following: PD-L1 + PD-1, PD-L1 + IDO, Lag3 + PD-1, and Lag3 + IDO. These pairs would be tallied in the overall interaction matrix for each patient, in which the value at row A and column B represents the number of times a cell positive for protein A was adjacent to a cell positive for protein B. For this analysis, we only counted interactions between functional proteins, excluding proteins used for lineage assignment.

We selected the top half of the symmetric matrix and flattened it to create feature vectors for each patient. Hierarchical clustering was performed and the dendrogram was cut to produce two clusters based on silhouette score analysis. These two clusters were compared using Kaplan-Meier curves, a two-sided log-rank test, and Cox regression. To assess the importance of individual interactions, we fit a random forest with all interactions as predictors and the cluster assignment as the response. We measured variable importance using the mean decrease in the Gini index.

### Healthy Tissue Analysis

We applied our computational pipeline to a set of 8 MIBI images of healthy tissue to validate our methods. We examined 6 proteins: FoxP3, IDO, Ki67, PD-1, PD-L1, and phospho-S6. We calculated the expression levels of the proteins in an identical fashion to the previous analysis of the TNBC images. We performed a two-sided Wilcoxon rank-sum test to compare expression levels between TNBC tissue and healthy tissue.

We also profiled cell-to-cell interactions in the healthy tissue in an identical manner to previous analysis. After obtaining interaction features, we performed dimensionality reduction using Uniform Manifold Approximation and Projection (UMAP)^49^. The resulting reduced features were compared to UMAP-reduced features of TNBC images.

### Multivariate Analysis

To assess whether the features identified by our computational pipeline contained independent prognostic information, we performed multivariate Cox regression. We fit three Cox Proportional Hazard models, each of which contained one of the three cluster variables identified by our study (protein co-expression, functional protein interactions, immunoregulatory protein interactions), two clinical variables (tumor grade and age), and the immune architecture distinction determined by Keren et al. We found the hazard ratio of each cluster variable and hypothesis tested the coefficient of each cluster variable to determine whether the variables contained additional prognostic information.

We also fit random forests to measure relative variable importance. We included all six variables in the random forests and calculated SHAP (Shapley Additive Explanations)^50^ values to get stable estimates of variable importance. SHAP values quantify the change in model prediction that would result from conditioning on a certain feature. They have been shown to be more aligned with human intuition regarding feature importance and attribution. We measured overall goodness-of-fit using Harrell’s c-index.

### Statistical Analysis

Primary statistical analyses were performed using Python (v3.7.3, Python Software Foundation, https://www.python.org/) with the lifelines (v0.24.0), scipy (v1.4.1), seaborn (v0.10.1), and pysurvival (v0.1.2) packages.

Pseudocode explaining the major steps of pipeline is shown in Supplementary Figure 7.

## Data Availability

MIBI images and other raw data for TNBC patients can be found at https://mibi-share.ionpath.com/. The link comes with an easy-to-use interface that allows for easy examination of the data upon registration. MIBI images for healthy patients will soon be made available on a Human Tumor Atlas Network public repository. The data produced by intermediary steps in the computational pipeline can be found at github.com/aalokpatwa/rasp-mibi/ in the intermediate_data/ folder.

## Code Availability

The computational pipeline used to produce the findings in this study can be found at github.com/aalokpatwa/rasp-mibi/blob/main/rasp_mibi_pipeline.ipynb. The pipeline is included in a Jupyter notebook with the output produced, as well as in separate .py files with instructions included.

Pseudocode describing the developed computational pipeline, including individual algorithms and techniques used, is shown in Supplementary Figure 7.

## Acknowledgments

This work was funded by the Stanford Departments of Pathology and Biomedical Data Science through a Stanford Clinical Data Science Fellowship to R.Y. The authors thank Biorender.com for their service in creating biological drawings.

## Author Information

### Affiliations

**Department of Biomedical Data Science, Stanford University, Stanford, CA 94305, USA** Aalok Patwa, Rikiya Yamashita, & Daniel L. Rubin.

**Archbishop Mitty High School, San Jose, CA 95129, USA**

Aalok Patwa

**Center for Artificial Intelligence in Medicine and Imaging, Stanford University, Stanford, CA 94305, USA**

Rikiya Yamashita, Jin Long & Daniel L. Rubin.

**Department of Pathology, Stanford University, Stanford, CA 94305, USA**

Michael Angelo

**Department of Molecular Cell Biology, Weizmann Institute of Science, Rehovot, Israel**

Leeat Keren

### Contributions

A.P. conducted the analysis and wrote the manuscript. R.Y. contributed to the design of the analysis substantially, providing constant feedback and interpretation of results. L.K. provided data, suggested analyses and evaluated results. J.L. suggested and reviewed analysis and results. M.A. provided data and interpreted results. D.R supervised the project. All authors participated in the critical revision and approval of the manuscript.

### Corresponding Author

Direct correspondence to Daniel Rubin.

## Ethics declarations

The authors declare no competing interests.

## Notes

https://mibi-share.ionpath.com

### Competing Interest Statement

The authors have declared no competing interest.

