## Supplementary Tables and Figures for "Multiplexed Imaging Analysis of the Tumor-Immune Microenvironment Reveals Predictors of Outcome in Triple-Negative Breast Cancer"

### Supplementary Information

| | |
|---|---|
| **Functional Proteins** | Beta Catenin, CD138, CD45RO, CD63, FoxP3, H3K27me3, H3K9ac, HLA-DR, HLA Class 1, IDO, Keratin17, Keratin6, Ki67, Lag3, p53, PD-L1, PD-1, Phospho-S6 |
| **Lineage Proteins** | CD11b, CD11c, CD16, CD20, CD209, CD3, CD31, CD4, CD45, CD56, CD68, CD8, dsDNA, EGFR, MPO, Pan-Keratin, SMA, Vimentin |

**Supplementary Table 1. Delineation between functional and lineage proteins**. Lineage proteins are those that are solely used for the identification of cell type, whereas functional proteins provide information related to proliferation or metabolic activity[25].

| Protein | Coefficient | Hazard Ratio | Coefficient P | BH-Corrected FDR |
|---|---|---|---|---|
| **CD45RO** | -0.019 | 0.981 | 0.051 | 0.311 |
| **PD1** | -0.112 | 0.894 | 0.063 | 0.311 |
| **CD138** | 0.018 | 1.018 | 0.069 | 0.311 |
| **IDO** | -0.067 | 0.935 | 0.084 | 0.311 |
| **HLA_Class_1** | -0.053 | 0.948 | 0.104 | 0.311 |
| **HLA-DR** | -0.013 | 0.987 | 0.12 | 0.311 |
| **Lag3** | -0.232 | 0.793 | 0.121 | 0.311 |
| **p53** | -0.023 | 0.978 | 0.212 | 0.477 |
| **H3K9ac** | 0.086 | 1.09 | 0.331 | 0.661 |
| **H3K27me3** | 0.724 | 2.063 | 0.367 | 0.661 |
| **Keratin17** | 0.009 | 1.009 | 0.432 | 0.684 |
| **Beta catenin** | -0.008 | 0.992 | 0.476 | 0.684 |
| **phospho-S6** | 0.012 | 1.012 | 0.494 | 0.684 |
| **Keratin6** | 0.005 | 1.005 | 0.651 | 0.837 |
| **PD-L1** | -0.004 | 0.996 | 0.715 | 0.858 |
| **CD63** | 0.003 | 1.003 | 0.791 | 0.890 |
| **Ki67** | -0.007 | 0.993 | 0.846 | 0.892 |
| **FoxP3** | -0.027 | 0.973 | 0.892 | 0.892 |

**Supplementary Table 2a. Protein expression Cox regression results for recurrence.** This table shows the results from performing univariate Cox regression with each protein's expression as predictors and recurrence outcome as the response. There is no association between protein expression and recurrence in the cohort.

| Protein | Coefficient | Hazard Ratio | Coefficient P | BH-Corrected FDR |
|---|---|---|---|---|
| **Keratin6** | 0.025 | 1.025 | 0.034 | 0.374 |
| **HLA-DR** | -0.018 | 0.982 | 0.045 | 0.374 |
| **Lag3** | -0.336 | 0.715 | 0.073 | 0.374 |
| **Keratin17** | 0.02 | 1.021 | 0.096 | 0.374 |
| **HLA_Class_1** | -0.055 | 0.946 | 0.109 | 0.374 |
| **H3K27me3** | -0.073 | 0.93 | 0.146 | 0.374 |
| **IDO** | -0.049 | 0.952 | 0.164 | 0.374 |
| **CD138** | 0.014 | 1.014 | 0.175 | 0.374 |
| **PD1** | -0.069 | 0.934 | 0.187 | 0.374 |
| **CD45RO** | -0.011 | 0.989 | 0.241 | 0.434 |
| **p53** | -0.014 | 0.987 | 0.353 | 0.578 |
| **CD63** | 0.008 | 1.008 | 0.511 | 0.671 |
| **Beta catenin** | -0.007 | 0.993 | 0.517 | 0.671 |
| **H3K9ac** | 0.056 | 1.058 | 0.522 | 0.671 |
| **PD-L1** | -0.006 | 0.994 | 0.626 | 0.751 |
| **FoxP3** | -0.085 | 0.918 | 0.688 | 0.774 |
| **Ki67** | 0.009 | 1.009 | 0.771 | 0.816 |
| **phospho-S6** | 0.001 | 1.001 | 0.928 | 0.928 |

**Supplementary Table 2b. Protein expression Cox regression results for survival.** This table shows the results from performing univariate Cox regression with each protein's expression as predictors and survival outcome as the response. There is no association between protein expression and survival in the cohort.

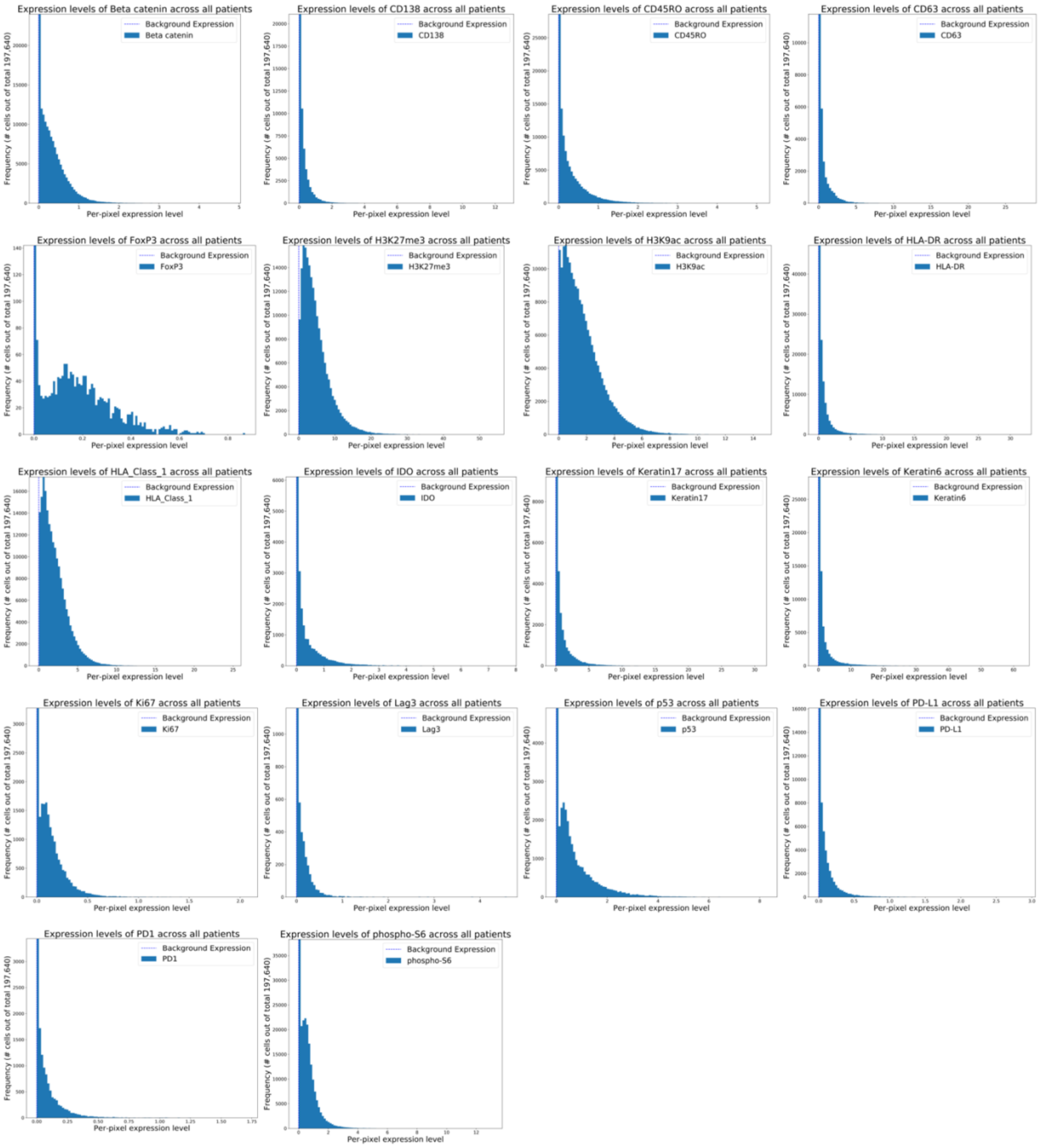


**Supplementary Figure 1. Histograms of expression for all functional proteins.** Each of the plots shown in the figure is a histogram of the per-pixel expression levels of that protein across all of the cells in all of the patients. The y-axis has been shrunk to better show the distribution, as in reality, the values are extremely right-skewed, and the frequency of 0 expression dwarfs the rest of the values. However, it is still valuable to see the rest of the distribution, as this is where the difference in cellular phenotype is made. As such, the y-axis upper limit has been made twice the second highest frequency.

| Features | Chosen Clusters | Mean Silhouette Score |
|---|---|---|
| Protein co-expression | 2 | 0.42 |
| | 3 | 0.18 |
| | 4 | 0.04 |
| | 5 | 0.04 |
| | 6 | 0.02 |
| Functional proteins interactions | 2 | 0.38 |
| | 3 | 0.11 |
| | 4 | 0.05 |
| | 5 | 0.04 |
| | 6 | -0.23 |
| Immunoregulatory proteins interactions | 2 | 0.47 |
| | 3 | 0.17 |
| | 4 | 0.06 |
| | 5 | 0.04 |
| | 6 | -0.29 |

**Supplementary Table 3. Silhouette score analysis on hierarchical clustering.** This table shows the mean silhouette score of hierarchical clustering based on the number of chosen clusters for each spatial analysis. In all cases, choosing two clusters led to the highest silhouette score.

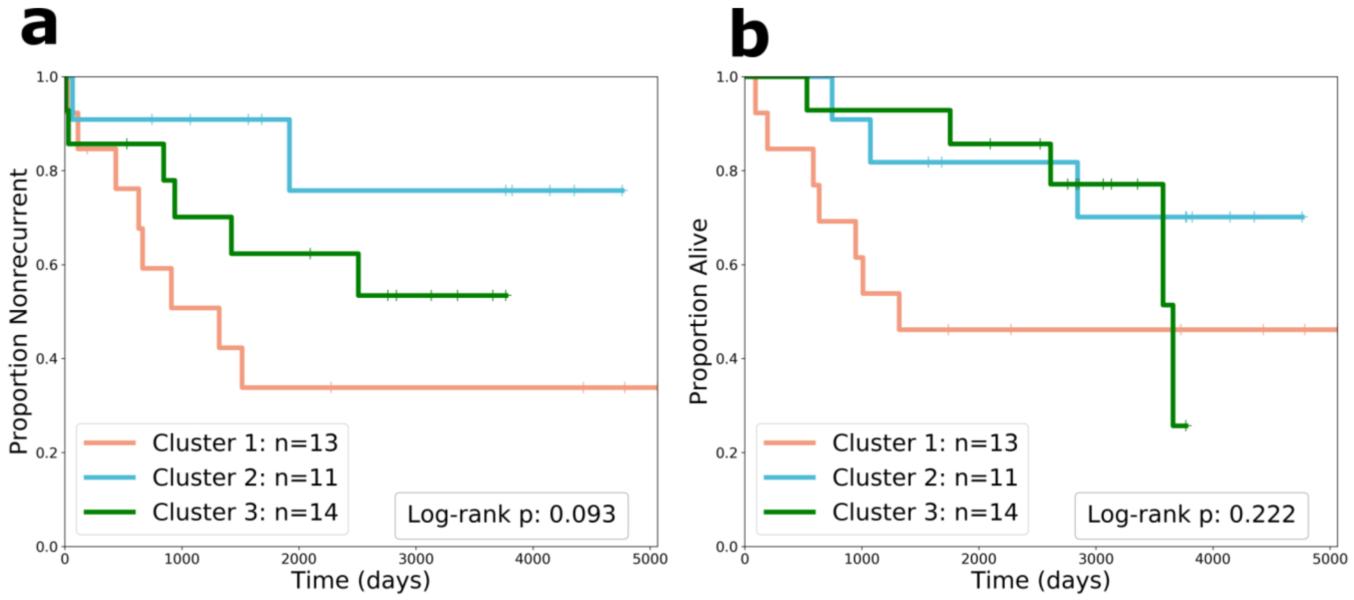


**Supplementary Figure 2. Kaplan-Meier curves comparing three patient clusters chosen from protein co-expression features. a** Kaplan-Meier curve comparing recurrence across the three patient clusters. The two-sided log-rank test p-value is shown. **b** Kaplan-Meier curve comparing survival across the three patient clusters. The two-sided log-rank test p-value is shown.

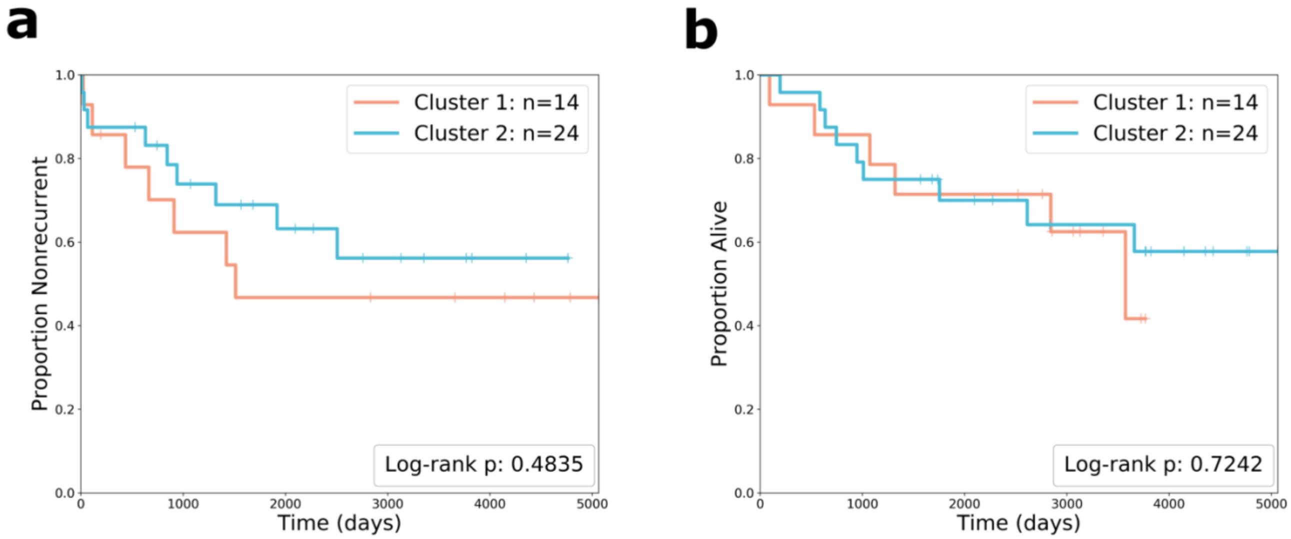


**Supplementary Figure 3. Kaplan-Meier curves comparing two patients clusters chosen from lineage proteins interactions features**. **a** Kaplan-Meier curve comparing recurrence outcomes across patient clusters formed from lineage proteins interactions features. There is not clear divergence between the clusters. The two-sided log-rank test p-value is shown. **b** Kaplan-Meier curve comparing overall survival across patient clusters formed from lineage proteins interactions features. There is no clear divergence.

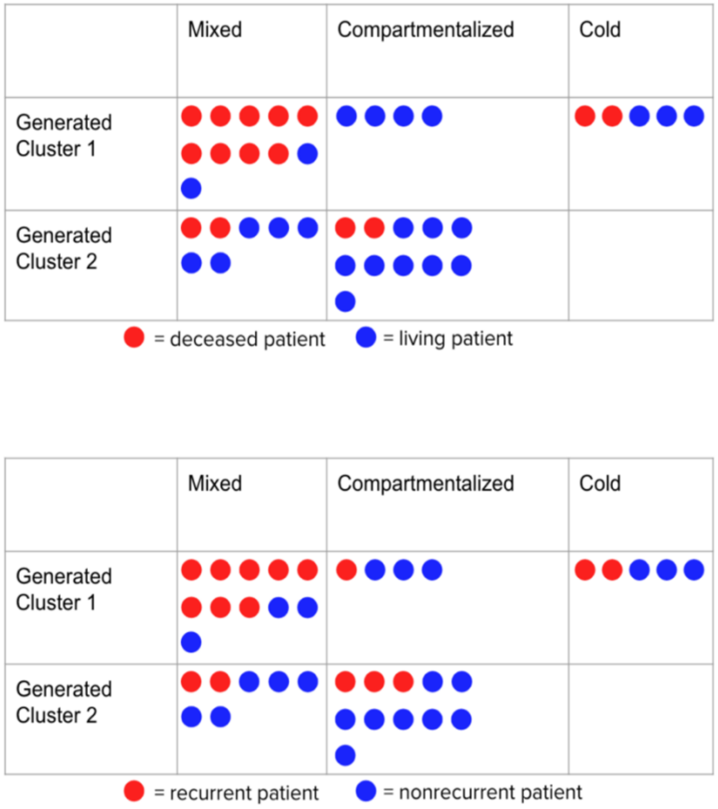


**Supplementary Figure 4.** Drawing comparing the patients in clusters formed from interaction features to Keren et al.'s designation of "mixed", "compartmentalized", and "cold" architectures.

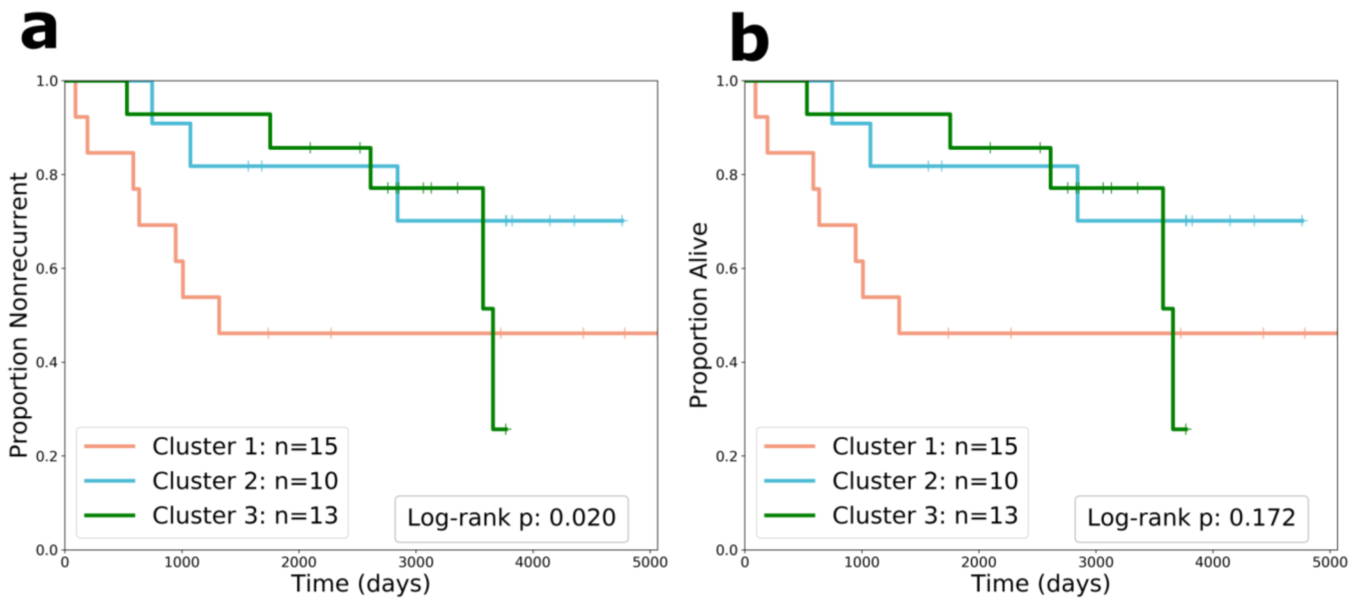


**Supplementary Figure 5. Kaplan-Meier curves comparing three patient clusters chosen from immunoregulatory protein interaction features. a** Kaplan-Meier curve comparing recurrence outcomes across the three patient clusters. The two-sided log-rank test p-value is shown. **b a** Kaplan-Meier curve comparing survival across the three patient clusters. The two-sided log-rank test p-value is shown.

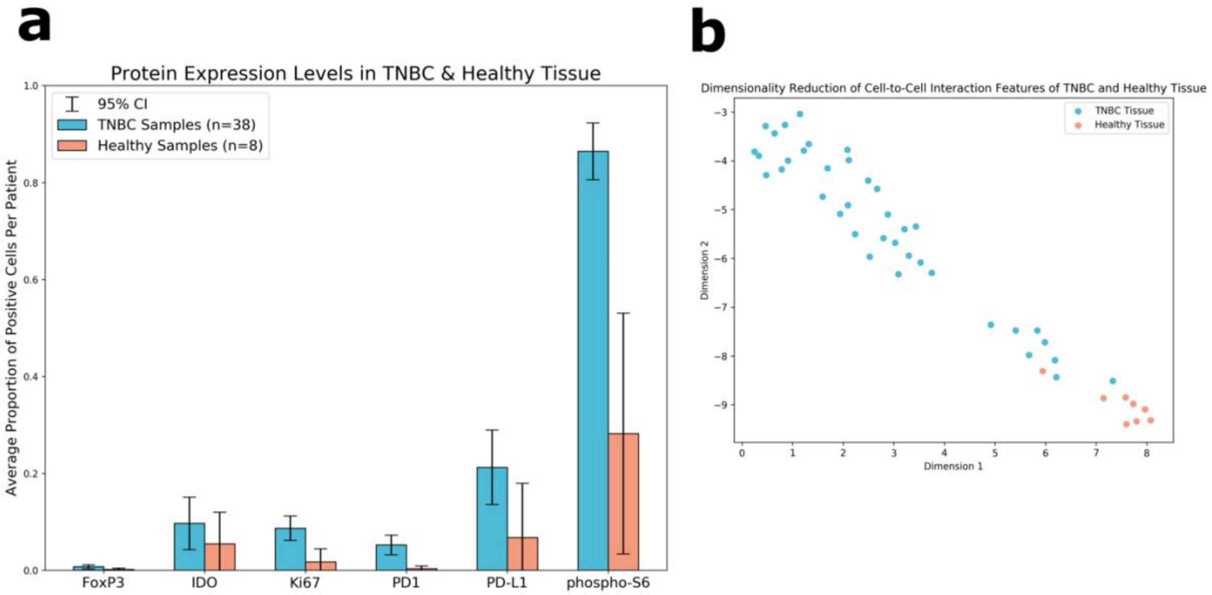


**Supplementary Figure 6. Healthy tissue analysis. a** Double bar plot comparing the expression levels of six proteins between TNBC samples and healthy samples. The error bars show the 95% confidence interval. **b** Scatterplot showing cell-to-cell interaction features reduced to two dimensions using UMAP. The healthy samples are bunched together in the bottom right corner.

```
Algorithm 1 RASP-MIBI Computational Pipeline
  input: image cohort of size N, segmentations of T cell types, MIBI images
  for K markers.

  # Immune Composition Analysis
  for cell type {c_t}_{t=1}^{T} do
    for patient segmentation {s_n}_{n=1}^{N} do
      numTotal ← total cells in s_n
      numCell ← number of cells of type c_t in s_n
      prevCell ← numCell / numTotal
    end for
    tTest(recurrence, prevCell)
    tTest(survival, prevCell)
  end for

  # Protein Expression Analysis
  for marker {m_k}_{k=1}^{K} do
    backgroundPixels ← total pixels of slide background across all patients
    backgroundExp ← \sum_{pixel=1}^{backgroundPixels} expression of m_k in pixel
    perPixelThreshold ← backgroundExp / backgroundPixels
    for patient MIBI image {p_n}_{n=1}^{N} do
      numTotal ← total cells in p_n
      numPositive ← 0
      for cell {C_j}_{j=1}^{numTotal} do
        numPixels ← number of pixels in C_j
        exp ← \sum_{pixel=1}^{numPixels} expression of m_k in pixel
        perPixelExp ← exp / numPixels
        positivityBin ← perPixelExp > perPixelThreshold
        if positivityBin then
          numPositive += 1
        end if
      end for
      proportionPositive ← numPositive / numTotal
    end for
    tTest(recurrence, proportionPositive)
    tTest(survival, proportionPositive)
  end for

  # Protein Co-Expression Analysis
  for patient MIBI image {p_n}_{n=1}^{N} do
    coexpMatrix ← matrix of co-expressions of protein pairs across all cells
  end for
  hierarchicalClustering(patients, coexpMatrix)
  logrankTest(cluster1, cluster2, recurrence)
  logrankTest(cluster1, cluster2, survival)
```

```
  # Voronoi Tessellation
  for patient MIBI image {p_n}_{n=1}^{N} do
    for marker {m_a}_{a=1}^{K} do
      for marker {m_b}_{b=1}^{K} do
        interactionMatrix[m_a, m_b] ← 0
      end for
    end for
    V ← drawVoronoiDiagram
    for edge in V do
      cell_p ← first cell in edge
      cell_q ← second cell in edge
      for (m_a, m_b) in interactionMatrix do
        if (cell_p positive for m_a AND cell_q positive for m_b) OR (cell_p positive
        for m_b AND cell_q positive for m_a) then
          interactionMatrix[m_a, m_b] += 1
        end if
      end for
    end for
    return interactionMatrix
  end for

  # Interaction Analysis
  hierarchicalClustering(patients, interactionMatrix)
  logrankTest(cluster1, cluster2, recurrence)
  logrankTest(cluster1, cluster2, survival)
```

**Supplementary Figure 7. Pseudocode for the main computational pipeline.** The major steps in the analysis pipeline are shown. Details regarding algorithms and techniques used are provided.
